# Biomechanical Compatibility With Physiological Limb-Flexion Deformation Reduces Inflammation and Neointimal Hyperplasia After Peripheral Stenting

**DOI:** 10.64898/2026.10.05.756779

**Authors:** Alexey Kamenskiy, Barbara Batista de Oliveira, Anastasia Desyatova, Frazer Heinis, Jordan Rosenbohm, Paul Deegan, Pranav Renavikar, Jason MacTaggart

## Abstract

**Introduction:** The femoropopliteal artery undergoes substantial axial compression, bending, torsion, and pinching during limb flexion. When an implanted stent does not accommodate these deformations, repetitive biomechanical mismatch may alter the vascular response to implantation and contribute to restenosis. We tested whether designing a peripheral stent to better accommodate physiologic limb-flexion deformation reduces the subsequent inflammatory and neointimal response.

**Methods:** A nitinol stent was developed using multi-objective computational optimization to maintain arterial apposition while reducing wall stress and luminal pinching during physiologic deformation. Sixteen Yucatan minipigs were studied in independent 3- and 6-month cohorts (n=8 each). Each animal received the optimized stent in one limb and a commercial stent with representative biomechanical behavior contralaterally. CTA quantified *in vivo* limb-flexion deformation, while μCT and histology assessed stenosis, neointimal accumulation, inflammatory-cell burden, αSMA-positive tissue, and extracellular-matrix composition.

**Results:** The optimized stent accommodated greater axial shortening than the commercial comparator at both 3 months (11.2±2.7% vs. 6.6±2.1%, p=0.003) and 6 months (13.7±4.4% vs. 6.8±2.3%, p=0.004). Intimal hyperplasia volume was lower with the optimized stent at 3 months (227.5±85.2 vs. 350.0±130.5 mm^3^, p=0.002) and 6 months (183.6±46.2 vs. 261.2±60.6 mm^3^, p=0.001), with significantly lower maximum, mean, and median stenosis at both timepoints. At 3 months, macrophage and total inflammatory-cell counts were 51% and 45% lower, respectively, with the optimized stent. Histologic tissue area was 28% lower at 3 months and 12% lower at 6 months. αSMA-positive tissue fraction was higher with the optimized stent at both timepoints, whereas elastin, glycosaminoglycan, and collagen fractions were similar between devices.

**Discussion:** Improved accommodation of limb-flexion deformation was associated with a smaller inflammatory and neointimal response to stent implantation through 6 months, while overall neointimal matrix composition remained largely unchanged. These findings provide *in vivo* evidence that biomechanical compatibility with the dynamic lower-limb environment can influence vascular healing and should be considered a mechanistic design criterion for peripheral artery stents.

## INTRODUCTION

Peripheral artery disease (PAD) frequently involves the femoropopliteal artery (FPA), where the durability of endovascular treatment remains limited despite advances in antiproliferative therapies^1^. Unlike more static vascular territories, the FPA undergoes large and spatially nonuniform changes in length and shape during routine limb flexion, including axial compression, bending, torsion, and cross-sectional distortion. These deformations are concentrated near the adductor hiatus and knee and can be further amplified by arterial calcification^2–5^. Current antiproliferative strategies target the downstream biological response to vascular injury but do not directly address this unusually demanding mechanical environment.

Implantation of a stent couples a metallic scaffold to an artery that normally changes length and shape with every limb movement. Commercial FPA stents differ substantially in their ability to accommodate axial compression, bending, and torsion^6^, and implanted stents modify native arterial deformation in a design-dependent manner^7^. Our prior work further demonstrated that stent-artery contact during cyclic limb deformation is dynamic and frictional, producing microscale abrasion, and that greater device abrasiveness is associated with poorer clinical patency^8^. Repetitive mechanical injury therefore provides a plausible biological link between biomechanical mismatch and restenosis by sustaining inflammation and neointimal growth.

These observations raise the possibility that a stent designed to accommodate physiologic limb-flexion deformation could alter the vascular response to implantation by reducing deformation-related mechanical mismatch. We previously used a multi-objective Design-of-Experiments framework to develop a nitinol FPA stent that maintains arterial apposition while reducing wall stress and luminal pinching under physiologic axial compression, bending, and torsion^9^. We subsequently established a porcine lower-extremity model that reproduces key geometric and deformation characteristics of the human FPA^10^. In the present paired study, we used this optimized design to test whether improved accommodation of limb-flexion deformation alters the vascular response to implantation. We therefore compared *in vivo* deformation, inflammatory-cell burden, α-SMA-positive tissue content, neointimal composition and accumulation, and stenosis at 3 and 6 months between the optimized stent and a commercial stent with representative biomechanical behavior.

## METHODS

### Animals and study design

All animal procedures were approved by the Institutional Animal Care and Use Committee of the University of Nebraska Medical Center and conducted in accordance with the U.S. Public Health Service Policy on Humane Care and Use of Laboratory Animals and the National Research Council’s Guide for the Care and Use of Laboratory Animals. Sixteen Yucatan minipigs were studied in independent 3- and 6-month cohorts (n=8 each). The 3-month cohort consisted of castrated males that were 9.8±0.4 months old and weighed 43.7±4.5 kg at baseline; the 6-month cohort consisted of females that were 8.2±0.4 months old and weighed 39.3±3.6 kg. Animals were maintained on a high-fat, high-fructose diet throughout the study, receiving 1 kg/day in two feedings. They received aspirin (650 mg/day) and clopidogrel (75 mg/day) beginning three days before stent implantation and continuing until sacrifice. One animal in the 3-month cohort was euthanized at 6.6 weeks because of a severe allergic skin reaction unrelated to the implanted devices. Bent-limb follow-up CTA was not obtained in this subject, but both stented arteries were included in postmortem μCT and histological analyses.

A paired, bilateral study design was used to isolate the effect of stent biomechanics. Each animal received the biomechanically optimized stent in one lower extremity and a commercial Zilver stent (Cook Medical) in the contralateral limb, with laterality randomized and balanced across animals. CTA was performed before implantation and again before sacrifice with the limbs in straight and flexed configurations to quantify arterial deformation. Following 3 or 6 months of implantation, stented arteries were harvested for μCT assessment of neointimal hyperplasia and stenosis followed by histological characterization of inflammatory-cell burden, tissue composition, and α-smooth muscle actin (αSMA)-positive tissue.

### Biomechanical stent design and comparator

The optimized stent was developed using a previously described multi-objective computational framework^9^. Briefly, finite-element simulations and Design-of-Experiments response-surface optimization were used to vary five geometric parameters while balancing arterial wall stress, stent-artery apposition, and luminal pinching under the combined axial compression, bending, and torsion experienced by the FPA during limb flexion. The resulting design consisted of sinusoidal nitinol rings connected by compliant sinusoidal links and was selected to accommodate physiologic deformation while minimizing adverse stent-artery interaction.

The optimized stent was manufactured from superelastic nitinol tubing with a nominal diameter of 6 mm and length of 40 mm using femtosecond laser cutting, shape-setting heat treatment, and electropolishing, and was crimped into a custom delivery system. A 6×40-mm bare-metal nitinol Zilver stent served as the mechanically contrasting commercial comparator. Zilver was selected because previous bench-top and perfused-cadaver studies from our group showed mechanical behavior representative of the range of commercial FPA stents under axial compression, bending, and torsion^6,7^.

### CTA and biomechanical analysis

CTA was performed before implantation and within one week before sacrifice using the straight- and flexed-limb configurations previously established for characterization of porcine lower-extremity biomechanics^10^. Animals were sedated with intramuscular tiletamine/zolazepam (4.4 mg/kg) and ketamine (2.2 mg/kg), intubated, and maintained under inhaled isoflurane (1-5% to effect). The straight posture was produced by extending and securing the hind limbs, whereas the flexed posture was obtained by flexing the limbs against the body during sternal recumbency.

CTA was acquired with a 256-slice GE Revolution CT scanner (GE Healthcare, Chicago, IL) at 120 kVp following intravenous administration of 100 mL iopamidol contrast. Images had a 512×512 matrix, 0.23-mm in-plane resolution, and 0.625-mm slice thickness. DICOM datasets were reconstructed in Materialise Mimics v25 (Materialise, Leuven, Belgium).

Arterial diameters were measured at the proximal, middle, and distal portions of the subsequently stented segment and in the major covered side branches. Limb-flexion-induced axial compression was calculated from the change in arterial segment length between straight and flexed configurations normalized to straight length. Tortuosity was defined as the difference between centerline length and end-to-end distance normalized to centerline length in the flexed configuration. Bending at the proximal and distal stent ends was quantified using inscribed-sphere radii and arterial angulation as previously described^10^.

### Stent implantation and follow-up

Stent implantation was performed under general anesthesia using percutaneous carotid access. Following placement of a 6-8 Fr sheath, the target FPAs were evaluated by intravascular ultrasound (IVUS; Philips Volcano, San Diego, CA). Six-millimeter stents were implanted in arteries with a target proximal diameter of ∼5 mm at peak systole. Animals received intravenous heparin (75 U/kg) before intervention.

Both stents were deployed distal to the inguinal ligament in the highly deformable proximal FPA segment previously shown to reproduce key features of the human adductor-hiatus mechanical environment^10^. In most animals, the proximal stent edge was positioned immediately proximal to the profunda femoris artery, the middle portion crossed the lateral circumflex femoral artery, and the distal edge terminated in the mid-Superficial Femoral Artery (SFA) (Figure 1A). Both devices were post-dilated with 6-mm angioplasty balloons using identical inflation protocols (8-10 atm for several seconds).

**Figure 1.**
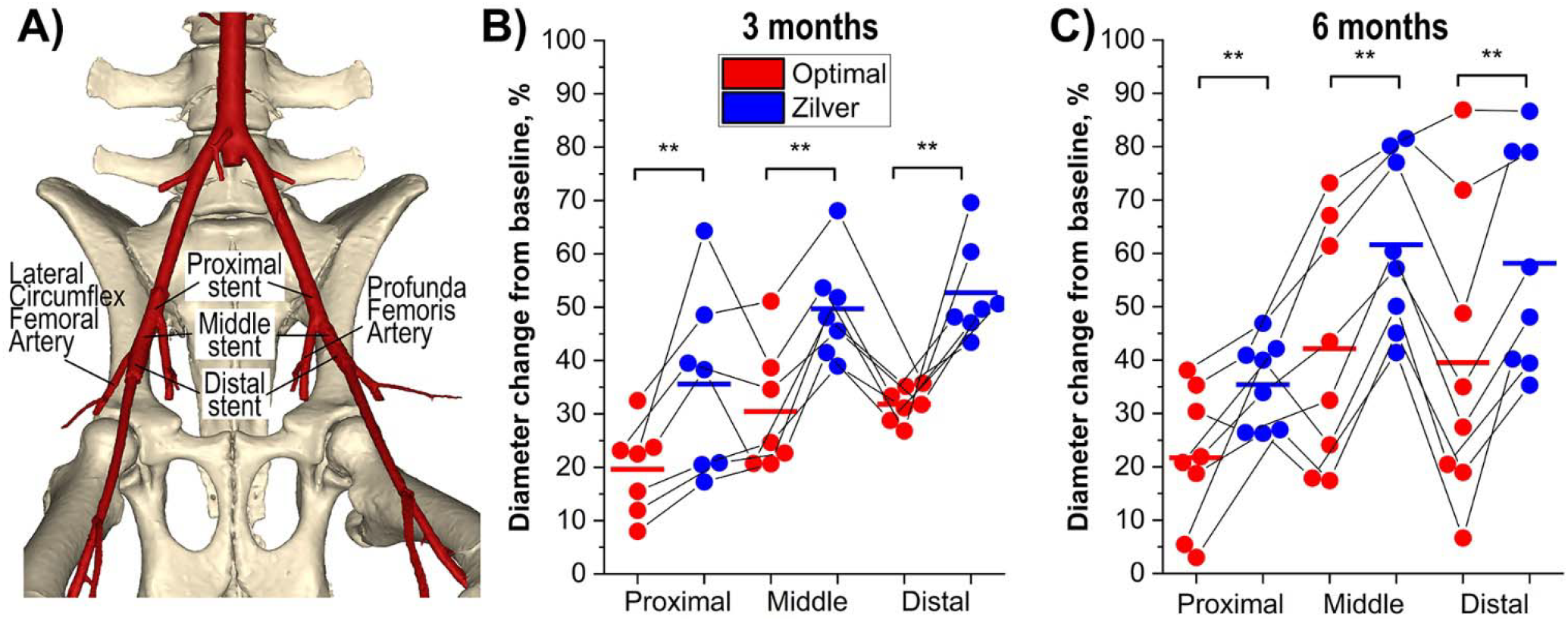
**A)** 3D CTA reconstruction showing the proximal, middle, and distal stent measurement locations and major arterial branches. **B-C)** Percent change in arterial diameter from baseline at 3 months **(B)** and 6 months **(C).** Paired Optimal and Zilver measurements from each animal are connected by lines; horizontal bars indicate group means. **p<0.01 for paired between-device differences in diameter change; n=7 at 3 months and n=8 at 6 months.

At study termination, animals were euthanized with intravenous pentobarbital (86.7 mg/kg) and phenytoin (11.1 mg/kg), followed by bilateral thoracotomy. Stented arterial segments were harvested with proximal-distal orientation preserved, flushed, branch vessels ligated, and pressure-fixed in formalin at 100 mmHg.

### μCT assessment of neointimal hyperplasia and stenosis

Pressure-fixed arteries were mounted on custom fixtures, protected from dehydration, and scanned using a μCT EasyTomS system (RX Solutions, Chavanod, France). Imaging was performed at 100-150 kV and 66-350 μA with four frames acquired per projection and a 0.35-mm copper filter to reduce stent-related beam-hardening artifacts. Isotropic voxel resolution was ∼30 μm. Reconstructed image stacks were imported into Materialise Mimics v25 for segmentation of the stent, arterial wall, and neointima. Neointimal volume was quantified by delineating tissue within the stent contour on cross-sectional images at the midline of the stent struts and subtracting the luminal volume. A centerline was fitted to the lumen, and cross-sectional lumen and stent-contour areas were calculated at 50 equally spaced axial locations spanning the entire stented segment. Percent stenosis was defined as the reduction in luminal area relative to the area enclosed by the stent contour at each location; maximum, mean, and median stenosis were calculated from the resulting stenosis profile over the full stented length.

### Histology

Following μCT, stented arteries were embedded in Spurr resin and sectioned at the proximal, middle, and distal stent regions using a contact-free femtosecond-laser sectioning system (TissueSurgeon; LLS ROWIAK, Hannover, Germany). Ten-micrometer sections were stained with hematoxylin and eosin (H&E) for inflammatory-cell analysis, Movat Pentachrome for elastin and glycosaminoglycans (GAGs), and Masson-Goldner Trichrome for collagen. Immunohistochemistry was performed to quantify αSMA-positive tissue as a surrogate for smooth muscle cell (SMC) content. Appropriate positive and negative controls were included. Because the 3- and 6-month specimens were processed separately, all Optimal and Zilver specimens within each cohort were sectioned and stained concurrently under identical conditions.

Slides were digitized at 40× magnification and analyzed in QuPath. For H&E sections, cell detection followed by a trained object classifier was used to distinguish macrophages/histiocytes from lymphocytes. Stain-specific pixel classifiers were used to quantify elastin-, GAG-, collagen-, and αSMA-positive areas. Stained areas were expressed both as absolute area and as a fraction of total tissue area. Analyses were performed at the proximal, middle, and distal stent locations.

### Statistical analysis

The animal was the experimental unit, and all within-cohort device comparisons accounted for the paired bilateral design. Normality of paired differences was assessed using the Shapiro-Wilk test. Device comparisons and baseline-to-follow-up changes were evaluated with paired t-tests when differences were normally distributed and Wilcoxon signed-rank tests otherwise. Differences in longitudinal change between devices were evaluated using paired change scores.

For histological analyses, measurements from the proximal, middle, and distal sections were aggregated within each artery before the primary device comparison. H&E inflammatory-cell counts were summed across the three sections, whereas absolute stained areas and area fractions were averaged. For location-specific analyses, paired device comparisons were performed separately at the proximal, middle, and distal locations, with Holm adjustment across the three locations within each endpoint.

Because the 3- and 6-month cohorts consisted of different animals, between-cohort comparisons were unpaired and performed using Welch’s t-test when both groups were normally distributed and the Mann-Whitney U test otherwise. Differences in device effect between cohorts were assessed by comparing the within-animal Optimal-Zilver differences between the 3- and 6-month groups using the same unpaired approach. Data are reported as mean ± standard deviation unless otherwise noted. All tests were two-sided, with p<0.05 considered significant.

## RESULTS

### Arterial remodeling

At baseline, arterial diameters at the proximal, middle, and distal stent locations did not differ between limbs assigned to either device in either cohort (all p≥0.121). After implantation, arteries enlarged with both devices, but the increase was consistently greater with Zilver. At 3 months, proximal, middle, and distal diameters increased by 19%, 30%, and 32% with the optimized stent and by 34%, 49%, and 52% with Zilver (all p≤0.018). Follow-up diameters were greater with Zilver at all three locations (all p<0.001; Figure 1B). At 6 months, corresponding increases were 21%, 39%, and 36% with the optimized stent and 35%, 60%, and 56% with Zilver (all p≤0.002), with Zilver-treated arteries again larger at all three locations (all p<0.001; Figure 1C). The Optimal-Zilver difference was greater in the 6-month than the 3-month cohort at the middle and distal stent locations (p=0.009 and p=0.023), but not proximally (p=0.218).

Profunda femoris artery diameter did not change significantly with either device at either timepoint. At 3 months, lateral circumflex femoral artery diameter increased by 22% with Zilver (p=0.017) but not with the optimized stent (p=0.194); neither device produced a significant change at 6 months. Direct Optimal-Zilver comparisons of either branch diameter were not significant at either timepoint (all p≥0.111).

### Accommodation of limb-flexion deformation

At baseline, SFA axial compression, tortuosity, and proximal and distal bending metrics did not differ between limbs assigned to either device in either cohort (all p≥0.130). Representative CTA images of the flexed limbs illustrate the different stent-artery deformation patterns produced by the optimized and Zilver devices in the same animal (Figure 2A,B). The optimized stent accommodated substantially greater axial shortening than Zilver at both 3 months (11.2±2.7% vs. 6.6±2.1%, p=0.003) and 6 months (13.7±4.4% vs. 6.8±2.3%, p=0.004; Figure 2C), with a similar device effect between cohorts (p=0.235). Overall SFA axial compression decreased significantly from baseline at 6 months with both devices (both p<0.001), but not at 3 months, and did not differ between devices at either follow-up (p=0.247 and p=0.548, respectively).

**Figure 2.**
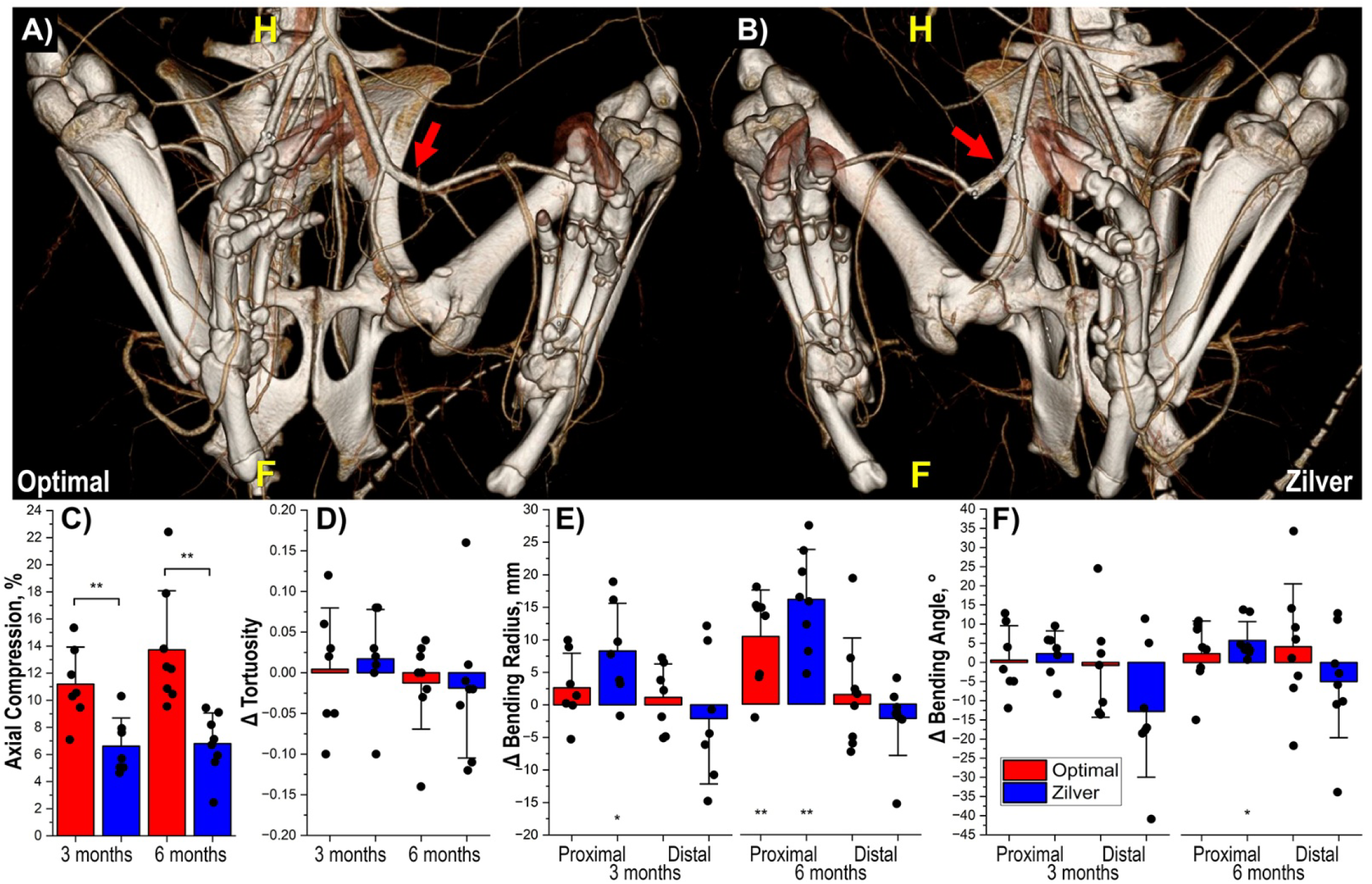
**A-B)** CTA of flexed limbs showing arterial geometry with the optimized **(A)** and Zilver **(B)** stents in the same animal. **C)** Stent axial compression at 3 and 6 months. **D-F)** Change from baseline in stent-region tortuosity **(D)**, proximal and distal bending radii **(E)**, and bending angles **(F).** Bars show mean ± SD with individual animals overlaid. Brackets indicate significant Optimal-Zilver differences; unbracketed asterisks indicate significant changes from baseline (*p<0.05, **p<0.01). n=7 at 3 months and n=8 at 6 months.

Stent-region tortuosity did not change significantly from baseline with either device and did not differ between devices at 3 months (p=0.709) or 6 months (p=0.182; Figure 2D). At 3 months, proximal and distal bending radii remained unchanged with the optimized stent, whereas Zilver increased proximal bending radius from 12.5±3.6 to 20.8±6.2 mm (p=0.024); direct device comparisons were not significant. At 6 months, proximal bending radius increased with both devices, without a significant between-device difference (p=0.243). Distal bending radius did not change significantly from baseline with either device, although the follow-up radius was greater with the optimized stent than with Zilver (13.7±7.2 vs. 8.0±2.2 mm, p=0.038; Figure 2E).

Bending angles showed no significant changes from baseline or between-device differences at 3 months. At 6 months, proximal bending angle increased with Zilver (144.9±4.7° to 150.7±4.3°, p=0.012) but not with the optimized stent (146.7±6.3° to 149.0±9.4°, p=0.469), although follow-up values did not differ between devices (p=0.607). Distal bending angles did not change significantly from baseline with either device, but the follow-up angle was greater with the optimized stent than with Zilver (134.8±14.8° vs. 121.5±9.3°, p=0.025; Figure 2F).

### Intimal hyperplasia and stenosis

Representative μCT reconstructions and cross-sections of contralateral optimized and Zilver stents from the same animal are shown in Figure 3A-D. At 3 months, maximum stenosis was lower with the optimized stent than with Zilver (45.4±8.7% vs. 56.7±12.7%, p=0.005), as were mean stenosis (30.3±8.8% vs. 41.5±11.8%, p=0.017) and median stenosis (29.8±8.9% vs. 41.7±12.4%, p=0.007; Figure 3E-G). Intimal hyperplasia volume was 1.54-fold lower with the optimized stent (227.5±85.2 vs. 350.0±130.5 mm^3^, p=0.002; Figure 3H).

**Figure 3.**
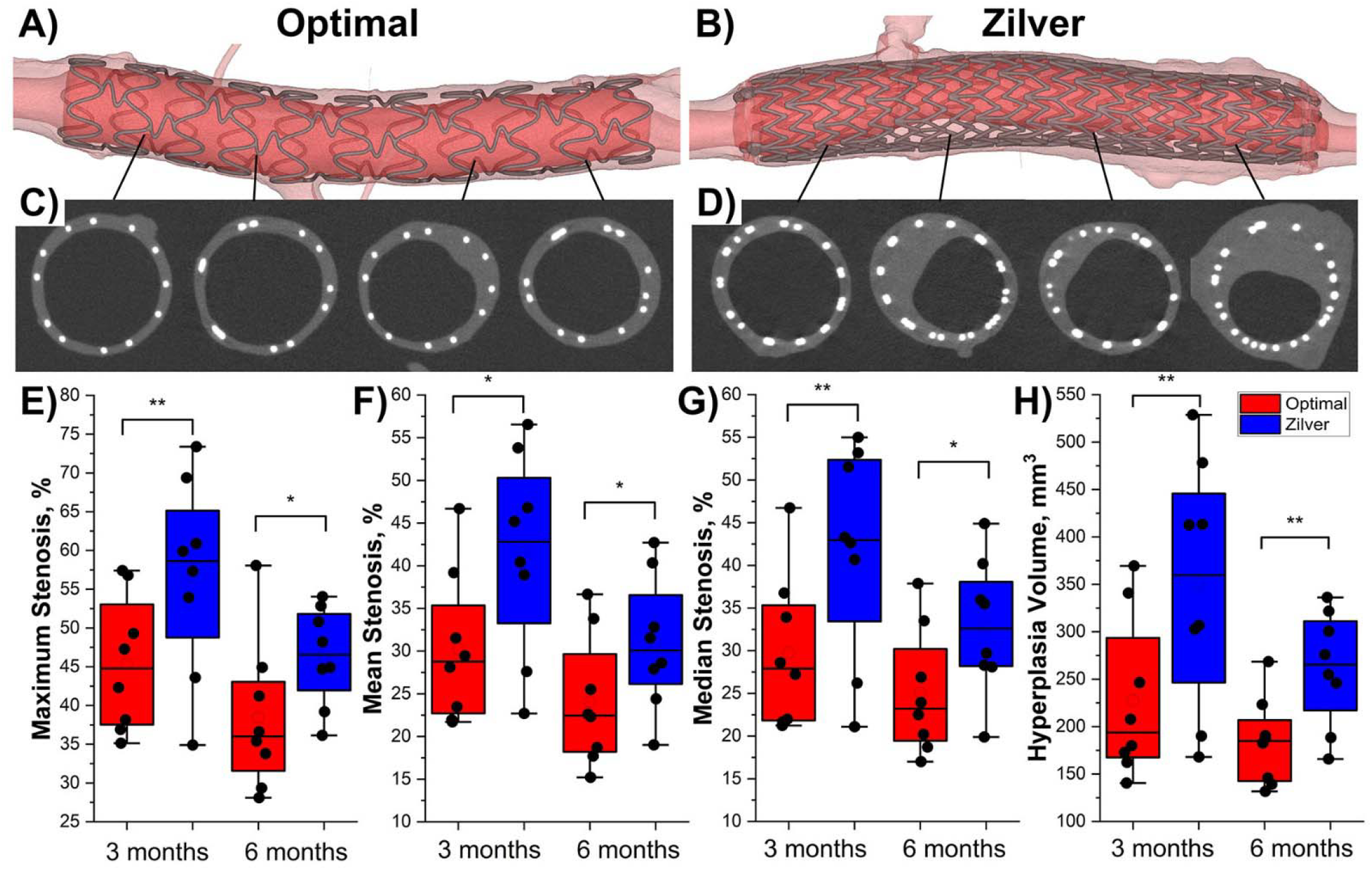
**A-D)** Representative μCT images of contralateral stented arteries from the same animal at 3 months. **A-B)** 3D reconstructions with the optimized **(A)** and Zilver **(B)** stents. **C-D)** Representative axial cross-sections showing the arterial wall, stent struts, and neointimal hyperplasia. **E-G)** Maximum **(E)**, mean **(F)**, and median **(G)** stenosis at 3 and 6 months. **H)** Intimal hyperplasia volume at 3 and 6 months. Boxes show the 25^th^-75th percentiles, center lines the median, hollow circles the mean, and black circles individual animals. *p<0.05, **p<0.01; n=8 at both timepoints.

The same pattern persisted at 6 months. Maximum, mean, and median stenosis remained lower with the optimized stent (38.4±9.7% vs. 46.4±6.4%, p=0.015; 24.1±7.6% vs. 30.9±7.8%, p=0.014; and 25.1±7.3% vs. 32.8±7.9%, p=0.010, respectively), and intimal hyperplasia volume was 1.42-fold lower (183.6±46.2 vs. 261.2±60.6 mm^3^, p=0.001). Zilver produced greater hyperplasia volume than the contralateral optimized stent in every animal at both timepoints. Neither stenosis nor hyperplasia differed significantly between the 3- and 6- month cohorts for either device (all p≥0.056), and the magnitude of the Optimal-Zilver difference was similar between cohorts (all p≥0.145).

### Histological response

Representative histological sections are shown in Figure 4. Consistent with the μCT findings, mean histologic tissue area was lower with the optimized stent at both 3 months (11.15±2.14 vs. 15.48±2.88 mm^2^/section, 28% lower; p<0.001) and 6 months (12.23±1.27 vs. 13.95±2.01 mm^2^/section, 12% lower; p=0.014; Figure 5F). At 3 months, this reduction was present across the proximal, middle, and distal stent regions, with 23-32% lower tissue area depending on stain and location, and remained significant after Holm adjustment. At 6 months, lower tissue area persisted in the middle (16% lower, adjusted p=0.044) and distal (20% lower, adjusted p=0.048) regions. Absolute collagen-positive area was also lower with the optimized stent at 3 months (1.01±0.25 vs. 1.72±0.70 mm^2^/section, p=0.015).

**Figure 4.**
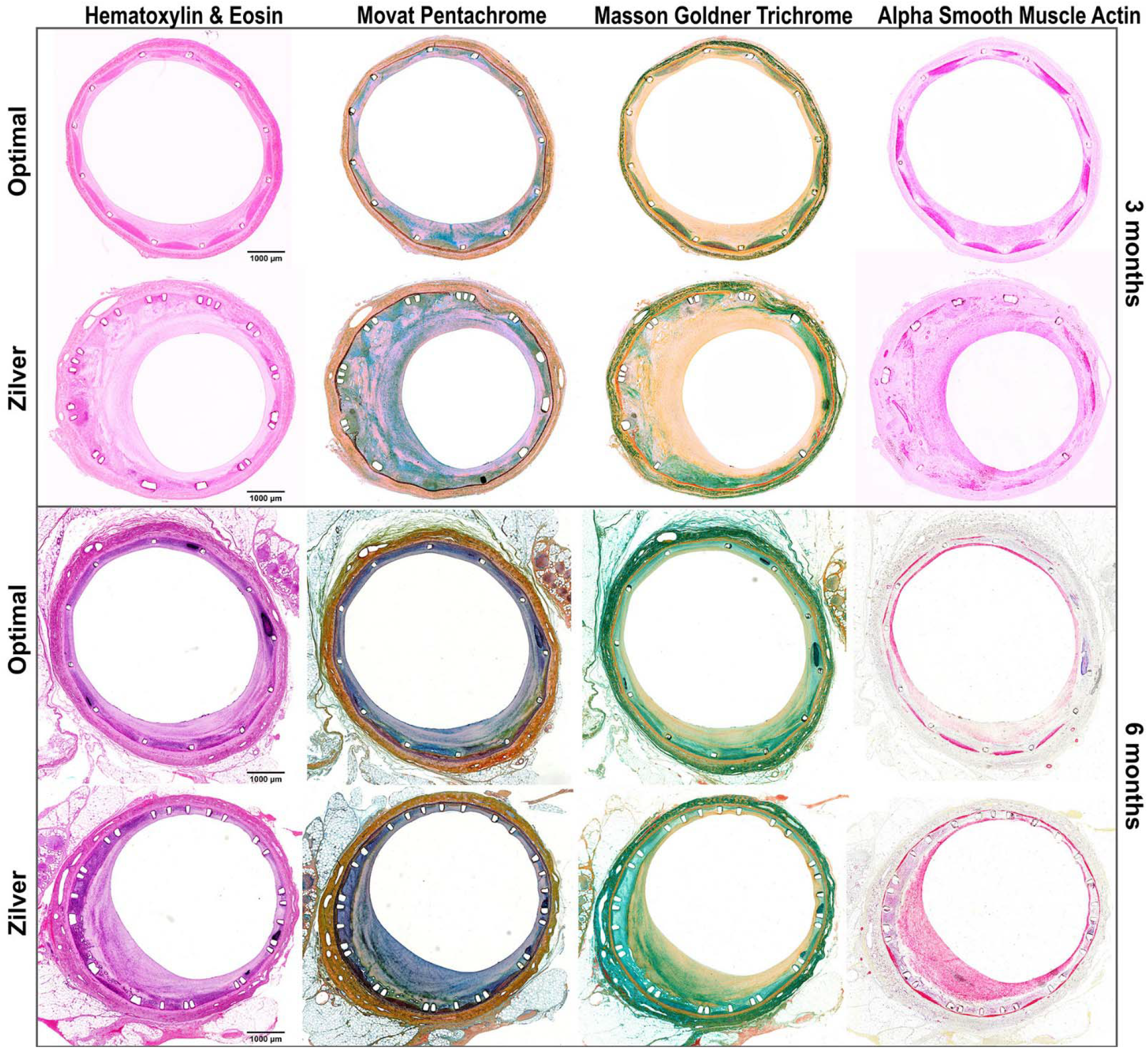
Representative histological cross-sections from the middle of contralateral stented arteries. Upper rows show the optimized and Zilver stents from the same 3-month animal; lower rows show the corresponding pair from a 6-month animal. Sections were stained with H&E, Movat Pentachrome, Masson-Goldner Trichrome, and αSMA to assess inflammatory cells, elastin/GAGs, collagen, and αSMA-positive tissue, respectively.

**Figure 5.**
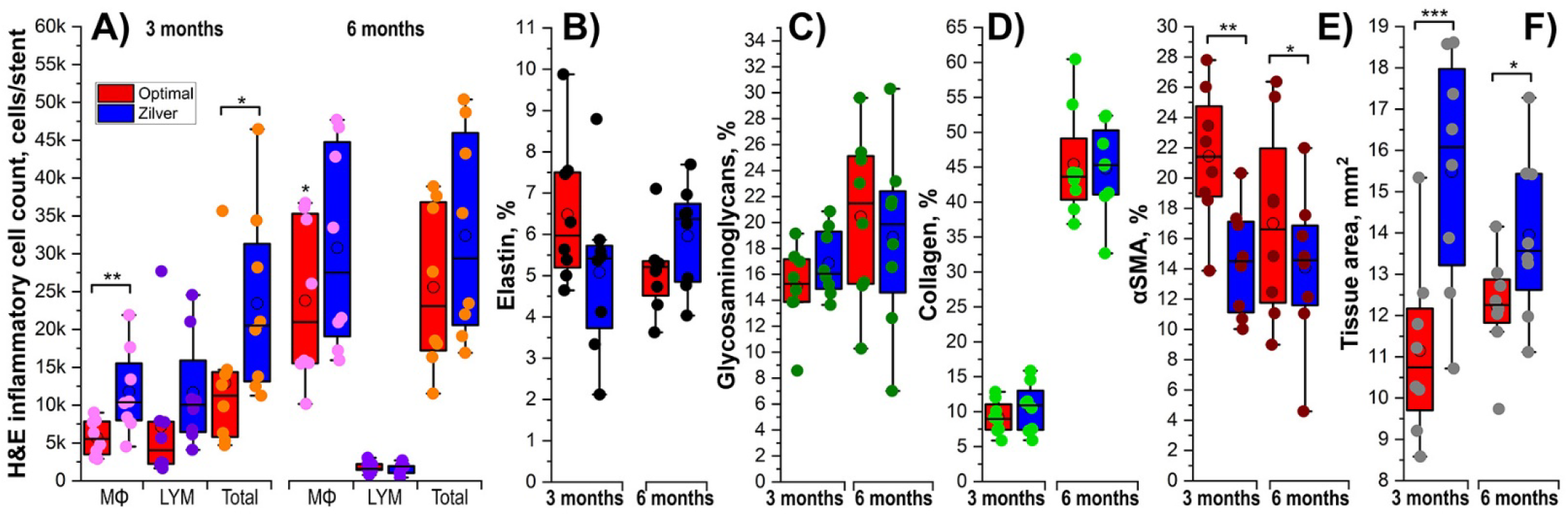
Quantitative histology of optimized- and Zilver-treated arteries at 3 and 6 months. **A)** H&E macrophage (MΦ), lymphocyte (LYM), and total inflammatory-cell counts summed across proximal, middle, and distal stent sections. **B-E)** elastin **(B)**, GAG **(C)**, collagen **(D)**, and αSMA **(E)** positive area as a percentage of total tissue area, averaged across the three locations. **F)** Mean histologic tissue cross-sectional area. Boxes show the 25^th^-75^th^ percentiles, center lines the median, hollow circles the mean, and points individual animals. Brackets indicate significant paired Optimal-Zilver differences (*p<0.05, **p<0.01, ***p<0.001); n=8 at both timepoints.

Inflammatory-cell burden was also reduced, particularly at 3 months. Macrophage counts were 51% lower with the optimized stent (5,711±2,366 vs. 11,772±5,670 cells/stent, p=0.002), and total macrophage-plus-lymphocyte counts were 45% lower (12,900±10,002 vs. 23,443±12,261 cells/stent, p=0.022; Figure 5A). Lymphocyte counts were numerically lower but not significantly different (p=0.123). At 6 months, macrophage and total inflammatory-cell counts remained 23% and 21% lower, respectively, but did not reach significance (p=0.093 and p=0.079). Distal macrophage counts at 3 months were 65% lower with the optimized stent (adjusted p=0.023).

Despite the difference in tissue accumulation, neointimal composition was largely similar between devices. Elastin, GAG, and collagen fractions did not differ significantly at either timepoint (all p≥0.108; Figure 5B-D). In contrast, the αSMA-positive fraction was higher with the optimized stent at both 3 months (21.4±4.5% vs. 14.5±3.6%, p=0.002) and 6 months (17.0±6.4% vs. 14.1±5.1%, p=0.025; Figure 5E), whereas absolute αSMA-positive area did not differ between devices. No location-specific differences in tissue composition remained significant after correction for multiple comparisons.

## DISCUSSION

This study tested whether designing a peripheral stent to better accommodate the large deformations imposed by limb flexion alters the biological response to implantation. The findings were consistent across independent mechanical and biological endpoints. The optimized stent accommodated approximately twice the axial shortening of Zilver at both 3 and 6 months, while arteries treated with the optimized device underwent less outward enlargement. These mechanical differences were accompanied by less neointimal accumulation and stenosis at both timepoints, lower inflammatory-cell burden at 3 months, and less histologic tissue accumulation. Despite this difference in tissue burden, extracellular-matrix composition was largely similar between devices, suggesting that improved biomechanical compatibility primarily reduced the magnitude of the vascular response rather than producing a fundamentally different type of neointimal tissue.

The different arterial diameter responses provide an additional indication that the two stents interacted differently with the vessel wall. Both devices had the same 6-mm nominal diameter and were implanted using the same sizing and post-dilation strategy, yet arteries containing Zilver enlarged more than those containing the optimized stent at the proximal, middle, and distal regions at both follow-up periods. Importantly, the optimized design was not intended simply to minimize radial force, but rather to maximize stent-artery apposition and minimize luminal pinching and arterial wall stress while accommodating the combined axial compression, bending, and torsion imported by limb flexion^9^. Differences in radial mechanical behavior between the devices, including chronic outward force and circumferential stiffness, may therefore contribute to the greater enlargement observed with Zilver. Once adequate apposition and luminal support are achieved, greater radial force is not necessarily beneficial and may instead increase chronic loading of the arterial wall. Experimental and clinical studies have linked stent-induced wall stress and sustained radial loading to arterial remodeling and intimal thickening^11,12^. Particularly relevant, Kawarada et al^13^ demonstrated that different bare nitinol platforms in the porcine FPA produced distinct patterns of vessel enlargement, injury, inflammation, and neointimal growth, supporting the concept that device mechanics can influence the subsequent vascular response.

The clearest biomechanical distinction between devices was their ability to accommodate axial shortening during limb flexion. Axial compression is a major component of physiologic FPA deformation, and restricting this shortening can redistribute deformation toward localized bending, kinking, pinching, and stent-end effects. The optimized stent accommodated substantially greater axial compression than Zilver at both follow-up periods, with a similar magnitude of device effect at 3 and 6 months. This extends our previous observations that commercial stents differ markedly in their mechanical response to lower-limb deformation^6^, that stent design alters FPA deformation after implantation^7^, and that cyclic stent-artery interaction is dynamic and frictional, with greater abrasiveness associated with poorer clinical patency^8^. Independent animal studies similarly show that vascular responses to nitinol stents depend on the mechanical environment: Sakaoka et al^14^ reported greater neointimal growth and persistent inflammation in dynamically loaded porcine FPAs than in less mobile femoral arteries, while Andrews et al^15^ observed greater neointimal hyperplasia in periarticular stents. These studies did not directly manipulate deformation accommodation as done here, but they independently support the premise that the mechanical environment surrounding a stent can influence the subsequent vascular response.

The bending measurements provided complementary, although less uniform, evidence of differences in local constraint. At 3 months, bending radii at both stent ends remained unchanged from baseline with the optimized device, whereas Zilver increased proximal bending radius, consistent with restricted deformation at the proximal stent edge. At 6 months, proximal bending radius increased with both devices, although the increase was numerically smaller with the optimized stent. Distally, neither device significantly changed bending radius from baseline; however, Zilver exhibited a smaller distal bending radius and sharper distal bending angle than the optimized stent at follow-up. Zilver also significantly increased proximal bending angle at 6 months, whereas the optimized stent did not. Although these sphere-based metrics describe overall arterial geometry rather than local contact mechanics and were not uniformly different between devices, their pattern is directionally consistent with the greater axial shortening accommodated by the optimized stent and the observed stent-end behavior.

Most importantly, the biomechanical differences were accompanied by a reproducible reduction in neointimal growth. At 3 months, intimal hyperplasia volume was 1.54-fold lower with the optimized stent, and maximum, mean, and median stenosis were all significantly reduced. The same pattern persisted at 6 months, when hyperplasia remained 1.42-fold lower and all three stenosis measures again favored the optimized stent. Zilver produced greater hyperplasia volume than the contralateral optimized stent in every animal at both follow-up periods, and the magnitude of the Optimal-Zilver difference was similar between cohorts. These findings extend prior evidence linking stent mechanics to neointimal response by prospectively altering deformation accommodation and demonstrating that the resulting mechanical behavior is associated with reduced tissue growth *in vivo*.

Absolute stenosis and hyperplasia values were numerically lower at 6 months than at 3 months for both devices. Because the two timepoints represent different animal cohorts, these data cannot establish longitudinal regression of neointima, and the between-cohort differences were not statistically significant. Prior porcine studies indicate that neointimal remodeling after nitinol implantation can vary with both implant location and device platform^13–15^. The more important observation here is that, regardless of any temporal remodeling, the relative advantage of the optimized stent remained evident at 6 months.

Histology provides further evidence that the mechanical differences were accompanied by a different biological response. At 3 months, macrophage burden was 51% lower and total inflammatory-cell burden was 45% lower with the optimized stent. At 6 months, macrophage and total inflammatory-cell counts remained numerically lower, although these differences were no longer statistically significant. Monocyte/macrophage recruitment is a central component of the vascular response to interventional injury and contributes to neointimal formation^16^. The 65% lower distal macrophage count with the optimized stent at 3 months is particularly interesting because stent-end regions are also where deformation transitions and local mechanical effects are most pronounced. Although tissue injury itself was not directly measured, the lower inflammatory-cell burden is consistent with reduced repetitive mechanical insult when physiologic deformation is better accommodated.

Histologic tissue area independently confirmed the μCT findings, being 28% lower with the optimized stent at 3 months and 12% lower at 6 months. At 3 months, reduced tissue accumulation was evident throughout the stented segment, whereas at 6 months significant differences persisted in the middle and distal regions. The agreement between volumetric μCT and section-based histology strengthens the conclusion that improved deformation accommodation was associated with genuinely less tissue accumulation rather than an effect specific to one measurement method.

Despite substantial differences in tissue accumulation, the composition of the neointima was largely similar between devices. Elastin, GAG, and collagen fractions did not differ significantly at either timepoint. Thus, improved biomechanical accommodation appears primarily to have reduced the amount of neointimal tissue formed rather than broadly changing its extracellular-matrix composition. One exception was αSMA-positive tissue fraction, which was higher with the optimized stent at both 3 and 6 months. Because absolute αSMA-positive area did not differ, this reflects a greater proportion of αSMA-positive tissue within a smaller neointima rather than greater absolute accumulation of such tissue. αSMA is associated with vascular SMC differentiation but is neither lineage-specific nor sufficient to establish phenotype^17^. Experimental stent studies have also shown that greater vascular injury can be accompanied by altered expression of SMC markers within the neointima^18^. The higher αSMA-positive fraction with the optimized stent is therefore compatible with a more organized or less injury-associated tissue response, but this interpretation remains tentative.

Taken together, the mechanical, μCT, and histological findings support a coherent mechanistic interpretation: a stent that better accommodates physiologic limb-flexion deformation produces less biomechanical mismatch with the artery, and this is associated with a smaller inflammatory and neointimal response to implantation. The present study does not directly measure microscopic tissue injury during limb motion or establish each intermediate step causally. However, the concordance among deformation accommodation, arterial remodeling, inflammatory-cell burden, tissue accumulation, and stenosis across two independent follow-up cohorts supports the concept that peripheral stent mechanics can influence vascular healing rather than merely determine device shape or structural performance.

Several limitations should be considered. The sample size was modest, although the paired contralateral design reduced inter-animal variability and the direction of the hyperplasia effect was highly consistent. The 3- and 6-month cohorts comprised different animals and differed by sex; comparisons between timepoints therefore cannot be interpreted as longitudinal changes, and sex may contribute to apparent temporal differences. Histological specimens from the two cohorts were processed in separate batches, although paired Optimal and Zilver samples within each cohort were processed concurrently, making within-timepoint comparisons the strongest histological inference. Only one commercial comparator was studied; Zilver was selected because previous work demonstrated biomechanical behavior representative of commercially available FPA stents^6,7^, but other architectures may produce different responses. Finally, implantation was performed in non-stenotic, non-calcified arteries, whereas clinical PAD frequently involves substantial calcification and altered wall mechanics^5,19^. Evaluation in disease-relevant models, including controlled peripheral arterial calcification^20^, will therefore be important to determine whether the same mechanobiological relationship persists in calcified PAD.

## CONCLUSIONS

This study demonstrates that the biomechanical compatibility of a peripheral artery stent with limb-flexion deformation can influence the vascular response to implantation. A stent designed to accommodate physiologic axial compression, bending, torsion, and pinching preserved greater arterial deformation and was associated with less inflammatory-cell accumulation, neointimal growth, and stenosis than a commercial comparator at both 3 and 6 months. Despite the marked difference in tissue burden, neointimal matrix composition was largely similar between devices, suggesting that improved deformation accommodation primarily reduced the magnitude rather than fundamentally altered the character of the healing response.

Together, these findings provide *in vivo* biological evidence that repetitive biomechanical mismatch is not merely a structural concern for peripheral stents but can contribute to vascular remodeling and restenosis. Incorporating physiologic limb-flexion biomechanics into stent design may therefore represent a mechanistically distinct strategy for improving the durability of femoropopliteal revascularization.

## ACKNOWLEDGEMENTS

This work was supported in part by the NIH awards HL125736, HL180371, and P20GM152301. The authors would like to thank the Tissue Analysis Core (TAC) of the NIH Center for Cardiovascular Research in Biomechanics (CRiB) for assistance with animal procedures.

## Notes

### Competing Interest Statement

The authors have declared no competing interest.

